# Multiple sclerosis: Effects of fixed-dose combination of dimethyl fumarate and NADPH oxidase inhibitor on oxidative stress markers and neurobehavioral activity in mice model of cuprizone-induced demyelination

**DOI:** 10.1101/2023.12.10.570981

**Authors:** Mani Gurubarath, Karthik Dhananjayan

## Abstract

**Background:** Motor coordination (MC) and long-term recognition memory (LRM) are affected in people with multiple sclerosis (MS). The exact mechanism underlying the pathogenesis of MS is still unknown. However, oxidative stress is one of the underlying mechanisms of the pathogenesis of MS that is detrimental to myelin due to an imbalance in the levels of antioxidants. Dimethyl fumarate (DMF) is a first-line treatment for relapsing-remitting multiple sclerosis (RRMS). Since OS plays a significant role in MS, in addition to DMF treatment, inhibition of NADPH oxidases (NOXs) may provide surplus glutathione (GSH) to prevent myelin loss. Hence, the objective of this study was to test the effect of a fixed dose of apocynin (50 mg/kg) combined with a fixed-dose DMF (30 mg/kg) on antioxidants, MC, and LRM in cuprizone (CUP)-induced mice model of demyelination.

**Methods:** The MC and LRM were evaluated by narrow beam and novel object recognition tests, catalase and superoxide dismutase activity by the degradation of hydrogen peroxide and nitroblue tetrazolium method; malondialdehyde by thiobarbituric acid assay, GSH by high-performance liquid chromatography, and presence of myelin by modified luxol fast blue staining.

**Results:** The combination therapy with DMF plus APO in CUP-fed mice preserved LRM and MC with a significant increase in catalase activity (1.15 ± 0.04 U/mg protein; p<0.001), superoxide dismutase activity (9.25 ± 0.10 U/mg protein; p<0.0001), and GSH (195.25 ± 22.75 nmol/mg protein, *\*\*\*\*p*<0.0001) of the cerebral cortex versus disease control (98.52 ± 8.85 nmol/mg protein). APO plus DMF30 in CUP-fed mice also preserved myelin (p<0.001) at the corpus callosum of mice brains.

**Conclusions:** We conclude that APO combination with DMF might have protected myelin by modulating (increasing) the levels of antioxidants to act against CUP-induced oxidative stress and thereby preserved MC and LRM in the cuprizone-induced mice model of demyelination.

## Background

Demyelination of neurons is a pathological process that leads to the destruction of the myelin lamellae with moderate parallel preservation of the axon and its supporting cells [1]. Demyelination slows down neurotransmission from the brain to various regions [2]. The function of neurons tends to deteriorate and cause problems depending upon the areas affected, including areas involved in memory and vision [3].

The demyelinating diseases of the central nervous system (CNS) are 1. multiple sclerosis (MS), 2. neuromyelitis optica, 3. acute disseminated encephalomyelitis, and 4. acute necrotizing hemorrhagic encephalomyelitis [4]. MS is one of the most occurring neurodegenerative, inflammatory, and autoimmune diseases of the central nervous system. It affects approximately 2.5 million people worldwide, particularly young adults between the ages of 20 and 40 [5]. The clinical patterns of MS are relapsing-remitting multiple sclerosis (RRMS), primary progressive multiple sclerosis (PPMS), secondary progressive multiple sclerosis (SPMM), and progressive relapsing multiple sclerosis (PRMS) [6]. People with any clinical pattern of MS experience disturbances in long-term recognition memory [7] and motor coordination [8].

The etiology of MS is still unclear. It is a multifactorial disease involving genetics, environmental factors [9], and various immune-mediated mechanisms [10]. At the molecular level, oxidative stress and inflammation play a role in the progress and promotion of the demyelination of the neurons[11]. The changes in the levels of lipids and the composition of lipids and fatty acids in the myelin are some of the general features of the demyelination process [12]. The decrease in the levels of anti-oxidants in myelin and axons causes uncontrolled lipid peroxidation [12]. Redox imbalance contributes to the oligodendrocyte susceptibility to oxidative stress-related damages [13]. Thus, it is essential to maintain surplus levels of anti-oxidants in the brain parenchyma.

Currently used drugs for MS are Glatiramer acetate, fingolimod, dimethyl fumarate (DMF), and laquinimod [14]. DMF is a drug of choice for first-line monotherapy in the early stages of MS [15]. DMF increases glutathione (GSH) levels *in vitro* and *in vivo*; however, it might not be sufficient to counteract the effect of oxidative stress. Thus, DMF combination with another drug may preserve and increase GSH in the brain parenchyma. Apocynin (APO), a NOX inhibitor, increases GSH levels in the cortex of rats and limits cell death following cerebral ischemia and post-spinal cord injury [16]. Systemic administration of APO reduces ROS, microglial activation, infiltration of immune cells, blood-brain barrier (BBB) disruption, and myelin loss in EAE-induced mice model of demyelination [17]. APO combination with DMF may protect the myelin sheath effectively against CNS demyelination.

Herein, we aimed to identify the effects of a fixed dose of APO in combination with a fixed dose of DMF on the levels of anti-oxidants and modulation of neurobehavioral deficits (motor coordination and recognition memory).

## Methods

### Chemicals

Cuprizone, Dimethyl fumarate, Apocynin, Malondialdehyde and Luxol fast blue (Merck, Germany). Cresyl violet, thiobarbituric acid (TBA), and reduced glutathione (Himedia, Mumbai); Tris (2-carboxyethyl)phosphine hydrochloride (TCEP) and 4-aminosulfonyl-7fluoro-2,1,3-benzoxadiazole (ABD-F) (TCI chemicals India Pvt. Limited, India). All solvents used for chromatographic analysis are of HPLC grade.

### Animals

The Institutional Animal Ethics Committee (158/PO/ReBi/Re-L/99/CPCSEA), PSG Institute of Medical Sciences and Research, Coimbatore, approved the study. Animals (C57BL/6J mice; sex: male; age: 8 weeks old, 24 ± 8 g, n=42 (7 mice per group)) procured from the PSG animal facility, PSG Institute of Medical Sciences and Research, Coimbatore. The mice were housed individually and maintained at a room temperature of 23 ± 01°C and humidity of 55 ± 05%, with 12 h of light & 12 h of dark. The study adheres to the guidelines of the Committee for the Purpose of Control and Supervision of Experiments on Animals (CPSCEA), New Delhi, India.

### Cuprizone (0.2%) mixed chow diet

The rodent chow pellets were powdered in a mixer grinder and weighed as per the daily needs of the mice. The diet contains chow pellets and cuprizone in a ratio of 99.8:0.2.

### Drug treatment

Mice were randomly divided into six groups, group I – Vehicle; group II - APO 50mg/kg plus DMF 30mg/kg; group III – Cuprizone (CUP) plus Vehicle; group IV – CUP plus APO 50 mg/kg, Group V - CUP plus DMF30 mg/kg; and Group VI - CUP plus APO 50 mg/kg plus DMF 30 mg/kg. Apocynin (APO) or dimethyl fumarate (DMF) was dissolved in mixture of propylene glycol-saline (4∶6, v/v) [18]. Mice (except group I and II) received 0.2% cuprizone mixed with a chow diet throughout the study (Days 01 to 35).

From day 11 to 35, mice in group I and III was treated with vehicle, whereas others (Group II, IV, V, and VI) were treated with corresponding drug or combination through oral gavage every morning between 8:30 a.m. to 9:30 a.m. The timeline (Fig.1) and dosage of the drugs were chosen as per the literature [19–21].

**Fig. 1.**
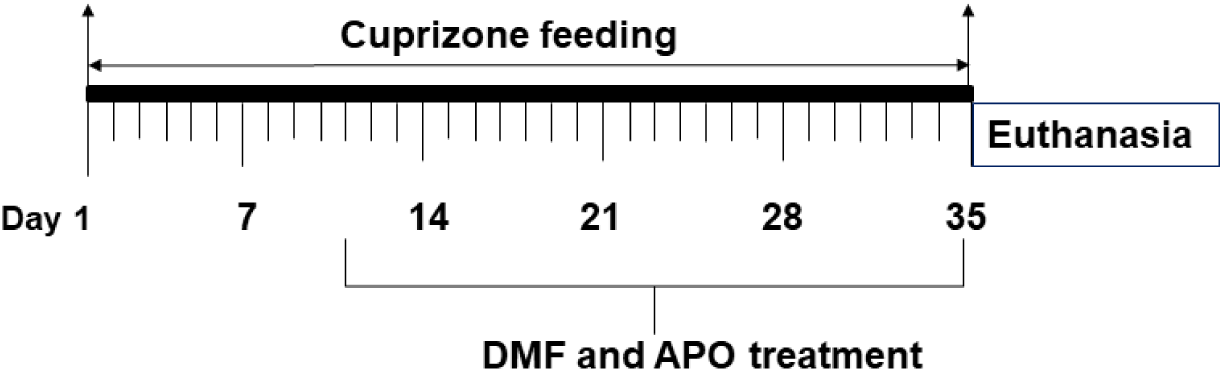
Study design.

### Behavioral studies

The narrow beam and novel object recognition tests were used to evaluate motor coordination and long-term recognition memory.

### Novel object recognition test

The novel object recognition test involves three phases (habituation, training, and testing). A square chamber contains equal quadrants with dimensions of 40 cm x 40 cm x 40 cm in length, breadth, and height, respectively. In the habituation phase, the mouse was placed at the center of an open, empty arena to explore for 10min. The next day (after 24h), in the training phase, the mouse was placed at the center and allowed to explore two identical objects kept in opposite quadrants for 10 min. After 24h (the testing phase), the mouse was placed at the center of the arena and allowed to explore one familiar (kept during the training phase) and a novel object in the opposite quadrant for 10 min. Note: At the end of the session, the mouse was removed from the arena and placed in a separate holding cage [22]. The total object exploration time, recognition, and discrimination index were calculated by the equation 1, 2, and 3, respectively.

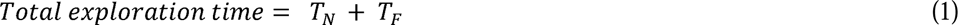

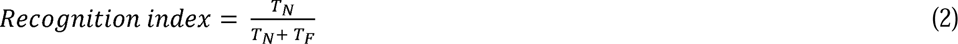

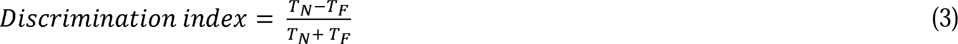

Where, T_N_-time spent with novel object; T_F_ – time spent with familiar object.

### Narrow beam test

The dimensions of the narrow beam used for this test is 100cm in length and 12mm in width and placed 60cm above the plane area whose ends consist of two enclosed safety boxes (a and b) (Supplementary information.1). The mouse was allowed to traverse a narrow beam from one point (c) to reach the other point (d) and then to an enclosed safety box. The time taken by the mouse (latency) to traverse the beam (80 cm) from one point to the other was noted (cut-off time: 60 sec). The test involves two consecutive trials, and the mean score of the two trials was calculated [23].

### Brain tissue isolation and preparation of homogenate

At the end of the study (on day 35), mice were euthanized by injecting xylazine (10 mg/kg, i.p.) and ketamine (40 mg/kg, i.p.) (Fig.1). Mice underwent transcardial perfusion by using phosphate-buffered saline (pH 7.4) [24]. Post perfusion, the brain was isolated, and transferred to corresponding beakers containing paraformaldehyde (4%) or stored at −80L. For biochemical analysis, the cerebral cortex including the corpus callosum was separated, weighed, homogenized with cold buffer (50 mM potassium phosphate, pH 7.4, 1mM EDTA, and 0.1% TritonX-100) and centrifuged at 1.6 x 10^4^ g for 15 min at 4L. After centrifugation, the supernatant was removed and stored at −80L. The amount of protein present in the supernatant was determined by Bradford assay (Supplementary information. 2).

### Determination of malondialdehyde

The malondialdehyde (MDA) in the samples was quantified by thiobarbituric acid assay [25]. The brain homogenate was diluted with phosphate-buffered saline (pH 7.4, 1:10). The diluted sample was mixed with 40 µL of sulfosalicylic acid (5%) and centrifuged at 1.6 x 10^4^ g for 20 min in an Eppendorf 5804 centrifuge. Post-centrifugation, 50 µ*L* of the sample was transferred to a 1.5 ml Eppendorf tube containing 200 µL of o-phosphoric acid (2%) and finally, 50 µ*L* of TBA was added, and incubated at 90L for 30 min. Post-incubation, 100 µL of the sample was transferred to a 96-well plate and the absorbance was measured at 532 nm (Thermo Scientific Multiskan GO UV/Vis microplate spectrophotometer). By using the linearity equation generated through the linearity curve, the amount of MDA present in the sample was determined and represented as per/mg of protein content.

### Measurement of glutathione

The measurement of glutathione was based on the HPLC-FLD detection of Thiol-ABD-F adduct obtained by pre-column derivatization of the sample with ABD-F at alkaline *p*H [26]. The sample was diluted with phosphate-buffered saline (pH 7.4, 1:10). The diluted sample was mixed with 40 µL of sulfosalicylic acid (5%) and centrifuged at 1.6 x 10^4^ g for 20 min in an Eppendorf 5804 centrifuge. Post centrifugation, a 50 µL of the supernatant was added to 1.5 ml Eppendorf tube containing 30 µL of a 1 mM tris(2-carboxyethyl) phosphine hydrochloride (TCEP) and incubated for 5 min at 35°C. Post-incubation, add 100 µL of borate buffer (0.1M, *p*H 9.3, with 1 mM EDTA) and 30 µL of the derivatizing agent ABD-F (1 mg/ml in 0.1M borate buffer *p*H 9.3 with 1 mM EDTA), incubate at 35°C for 10 min. Finally, the reaction was stopped by the addition of 50 µL of 2M hydrochloric acid. 10 µL of the derivatized sample was injected into the HPLC-FLD system. The HPLC-FLD system is composed of a Waters HPLC 515 binary pump, and a manual sample injector coupled with a 2475 fluorescence detector (Excitation:390 nm and Emission:510nm). A Waters C_18_ sun-fire column (5µm; 4.6 x 250 mm) was used for the separation of analytes. A mobile phase of acetate buffer (*p*H 9.3) and methanol (60:40) was used with a flow rate of 1 ml/min. The concentration of GSH (nmol/mg protein) present in the brain samples was determined from the linearity curve of the reference standard (Supplementary information. 3) and represented as per/mg protein content.

### The catalase and superoxide dismutase activity

The catalase activity was determined by the degradation of hydrogen peroxide [27]. A mixture was prepared by adding 100 µL of the sample to 900 ml of 0.25M phosphate buffer (*p*H 7.0). To this mixture, 1 mL of hydrogen peroxide (H_2_O_2_) solution was added and the absorbance was measured at 260 nm at room temperature for 30 sec. The results were expressed as CAT activity (U/mg of protein).

The superoxide dismutase (SOD) activity was measured by the nitroblue tetrazolium method [28]. A hundred microliter of the sample was added to a mixture containing 1.2 ml of sodium pyrophosphate buffer (pH 8.3; 0.052 M) and 300 µL of nitro blue tetrazolium (300 μM). Finally, 200 µL of NADH (750 μM) was added and immediately incubated at 30 °C for 90 sec. Post-incubation, the reaction was stopped by the addition of 100 µL of glacial acetic acid; after one minute 4.0 ml of nLbutanol was added and the reaction mixture was stirred. The color intensity of the sample was measured spectrophotometrically at 560 nm and the concentration of SOD was expressed as U/mg of protein.

### Histopathology (Modified luxol fast blue staining)

The isolated brains were post-fixed in paraformaldehyde (4%) for 48h at room temperature. The coronal section of mice brain was transferred to labelled cassettes, processed, and embedded in paraffin according to standard embedding techniques (using a Leica EG1150H paraffin embedding station and Leica EG1150C cold plate). The embedded tissue blocks were carefully sliced into 15µm sections using Leica RM2235 manual rotary microtome and mounted to glass slides. The sections were first hydrated in double distilled water (DDW) and rinsed in 70% ethanol and placed in 0.1% Luxol fast blue (Sigma) staining solution for 6h at 60 °C. After staining, the sections were then rinsed in 95% ethanol and DDW. Next, the sections were post-fixed in 1% lithium carbonate until there was a clear differentiation between white and grey matter staining. Sections were then counter-stained in 0.1% cresyl violet solution (Sigma) and then dehydrated in an ascending series of ethanol, treated with xylene, and protected with coverslip using a DPX mounting medium [29]. The stained sections were visualized using a Nikon H66L phase contrast microscope equipped with a digital camera (Nikon DS-Fi3 camera) and whole-slide digital images were acquired using an imaging system (NIS-Elements D 4.60.00 build 1171). The corpus callosum area of the images were analysed using ImageJ software and the extent of the myelination was scored using a scale ranging from 0 - Complete demyelination, 1 - very low myelination, 2 - low myelination, 3 - intermediate myelination, to 4 - Complete myelination.

### Statistical Analysis

The data were analyzed using GraphPad Prism Software, Inc., La Jolla, CA, USA version 8.02. The significant differences in the levels of biochemicals and enzyme activity were analyzed by one-way ANOVA, followed by Dunnett’s Post-hoc test. For behavioral assays, the significance in the groups was analysed by two-way ANOVA, followed by Tukey’s multiple comparison test.

## Results

Mice (group III to VI) were fed with 0.2% cuprizone and treated with a corresponding drug or combination or vehicle every morning (8:30 to 9:30 am). Analysis indicated a significant difference in the body weight among the groups fed with cuprizone during the study (F (24, 70) = 1.796; *p<0.05). Multiple comparison tests indicated a significant decrease in body weight of disease control (CUP plus Vehicle) on day 35 (5th week) (p<0.01) versus day 01. There were fewer weight changes in treatment groups, but not significant; the weight changes shown here are in the order of decrease in weight: Disease control < CUP plus APO50 < CUP plus DMF30 < CUP plus DMF30 plus APO50 < APO50 plus DMF30 or normal control.

### Effect of apocynin combination with DMF on long-term recognition memory in cuprizone-fed mice

The analysis showed no significant changes in TET of mice among the groups (F (5, 60) = 0.7889, p=0.5618). At baseline, all the mice spent more time in exploring the novel object without any significant differences. However, on day 35, there was a 0.5-fold decrease in total exploration time (TET) of disease control (p=0.3268) versus baseline with no significant changes (Fig.3A).

Next, analysis of RI indicated significant differences between the groups (F (5, 60) = 7.301, p<0.0001). On day 35, no significant changes in RI of cuprizone unfed, fixed dose combination (APO50 plus DMF30) treated mice (p>0.99) and vehicle treated normal control mice (p>0.99) versus their corresponding baseline values (day 01). In disease control, the RI decreased (0.5-fold) significantly on day 35 (p <0.0001) versus baseline (day 01). In CUP plus APO50 treated mice, there was only 0.83-fold decrease in RI but it was not significant (p= 0.6383) versus baseline, which indicates that the recognition memory was slightly preserved (Table 1). Mice treated with CUP plus DMF30 showed 0.7-fold, significant decrease (p=0.011) in the RI on day35 versus baseline (day 01). In cuprizone fed, fixed dose combination group (CUP plus APO50 plus DMF30), no significant changes in the RI (p>0.99) versus baseline (Fig. 3B).

Analysis of DI also indicated significant differences in the groups (F (5, 60) = 10.49, p<0.0001). The DI was unaffected in vehicle treated normal control mice (p>0.99) and cuprizone unfed, fixed dose combination (APO50 plus DMF30) treated mice (p>0.99) versus their corresponding baseline (day 01). In disease control, the discrimination index significantly decreased (2.41-fold) on day 35 (p<0.0001) versus day 01. In CUP plus APO50 treated mice, insignificant, 0.76-fold decrease in RI (p>0.99) versus baseline. Next, CUP plus DMF30 also showed an insignificant, 0.43-fold decrease in DI (p>0.99) versus baseline. In cuprizone fed, fixed dose combination group (CUP plus APO50 plus DMF30), the DI was unaffected and no significant changes compared to their baseline (p>0.99) (Fig.2C)

**Fig. 2.**
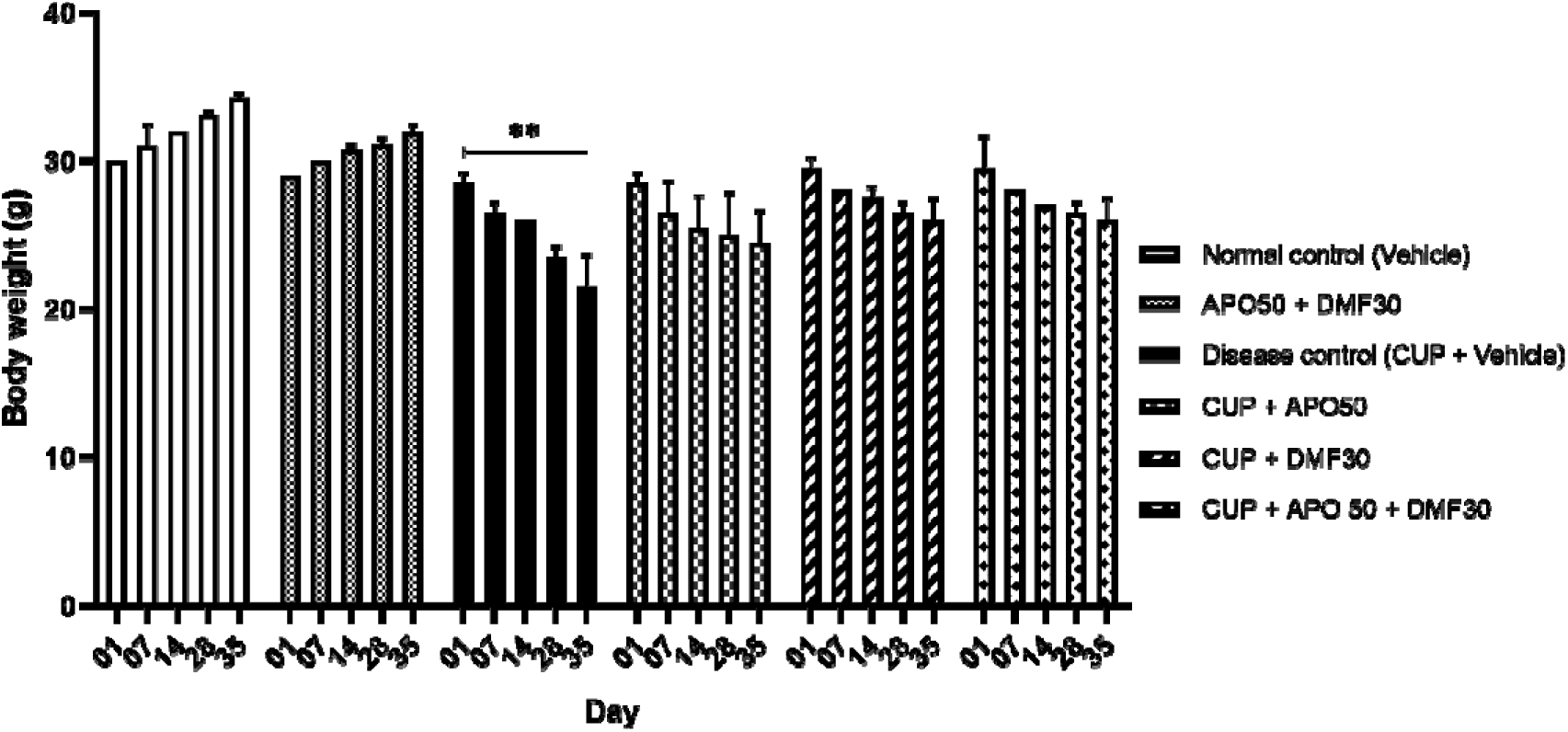
Body weight. The body weight of mice (mean ± S.D) plotted on y-axis against days on x-axis. The significance in weight of the mice between the days are represented by **p<0.01, n= (6 x 7 = 42).

### Effect of apocynin combination with DMF on motor-coordination in cuprizone-fed mice

The analysis of data indicated significant difference among the groups on the latency to traverse the distance between the beam (F (15, 120) = 3.144, p<0.001). No significant changes in latency to traverse between the two-points of the beam by cuprizone unfed, fixed dose combination (APO50 plus DMF30) treated mice (p>0.99) and vehicle treated normal control mice (p>0.99) versus their corresponding baseline (Table.2). In cuprizone fed disease control rats, the latency to traverse the distance increased significantly to 2.06- and 2.90-fold on days 28 (p=0.0263), and 35 (p<0.0001) respectively, compared to their baseline (Table.2).

The latency to traverse the distance decreased insignificantly to 0.94-fold by CUP plus APO50 treated mice on days 14 (p>0.99), however the latency increased insignificantly to 1.15- and 1.22-fold on days 28 (p>0.99) and 35 (p>0.99), respectively compared to baseline (Table.2). In CUP plus DMF30 treated mice, the latency increased insignificantly to 1.23-, 1.17-, and 1.18-fold on days 14 (p=0.97), 28 (p=0.99) and 35 (p=0.99) versus baseline (Table.2). In cuprizone fed, fixed dose combination group (CUP plus APO50 plus DMF30), the latency increased only 1.13-, 1.18- and 1.33-fold with no significant changes on day −14 (p>0.99), −28 (p=0.99) and −35 (p>0.99) compared to their baseline (Table.2) (Fig.3).

**Fig. 3.**
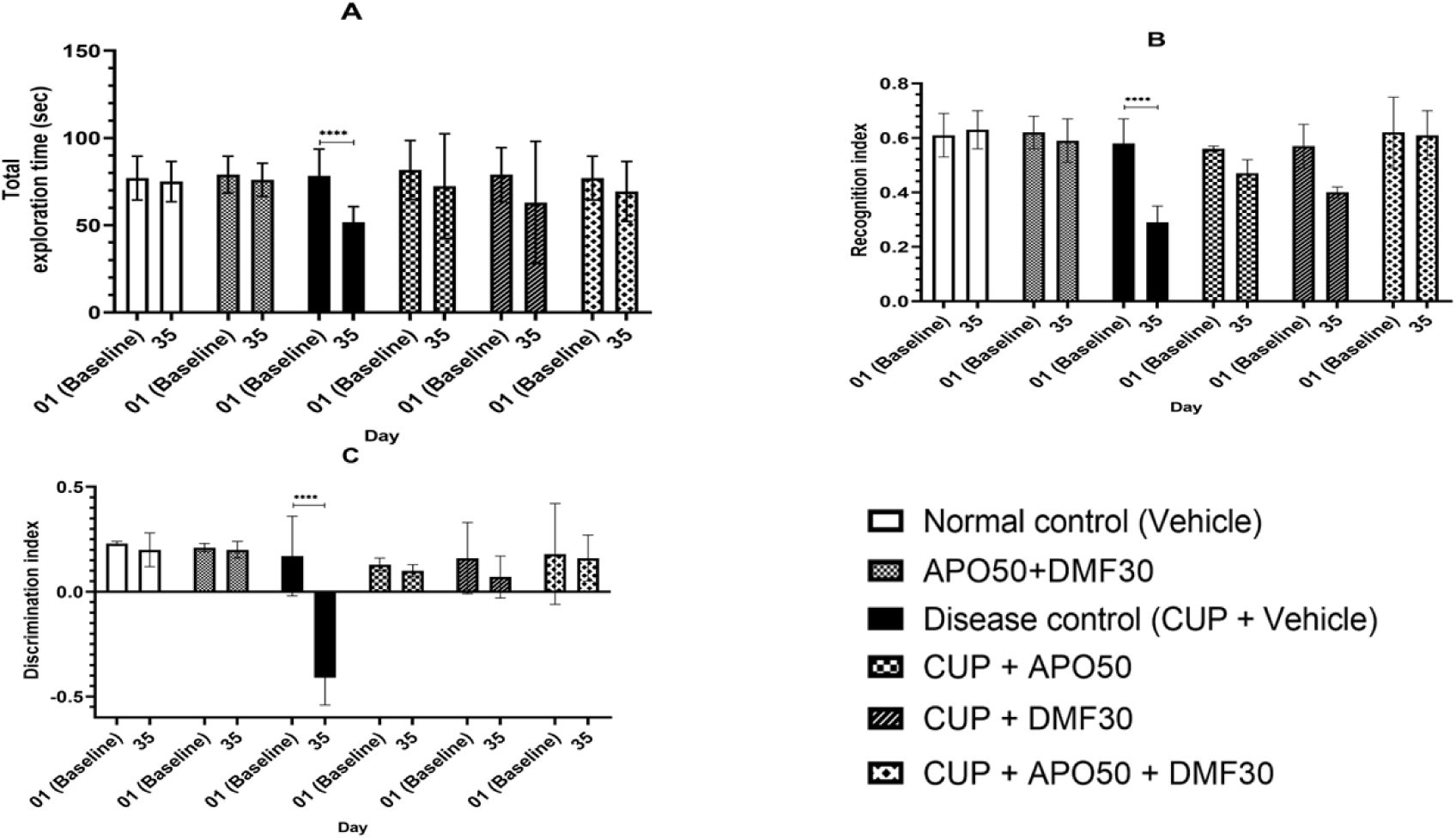
Novel object recognition test. (A) Total exploration time (TET): The TET (sec) was plotted on the y-axis against the day on the x-axis. (B) Recognition index (RI): The ability of the mice to recognize familiar and novel objects; the RI of the groups are plotted on the y-axis against day on the x-axis (C) Discrimination index (DI): The ability of the mice to discriminate between familiar and novel object; DI of the groups are plotted on the y-axis against day on the x-axis. All values are represented as mean ± SD and **** indicates p<0.0001, n=6×7= 42.

### Effect of APO combination with DMF on lipid peroxidation

The treatment with APO plus DMF30 showed significant differences in the MDA levels in the CCO (cerebral cortex including corpus callosum) of brain (F (5, 24) = 13.67, p<0.0001). MDA levels significantly increased in CCO of disease control (13.57 ± 0.25 nmol/mg protein; p<0.0001) versus normal control. In cuprizone unfed, APO50 plus DMF30, the MDA levels in CCO of brain was significantly less (7.98 ± 2.92 nmol/mg protein; p<0.0001) compared to disease control. In CUP plus APO50 (10.86 ± 0.43 nmol/mg protein; p=0.0269) and CUP plus DMF30 (9.75 ± 0.33 nmol/mg protein; p=0.0015), MDA levels significantly decreased in CCO of brain versus disease control. In CUP plus APO50 plus DMF30, MDA significantly reduced in CCO of the brain (7.92 ± 0.36 nmol/mg protein; p<0.0001) compared to disease control (Fig. 4A).

**Fig. 4.**
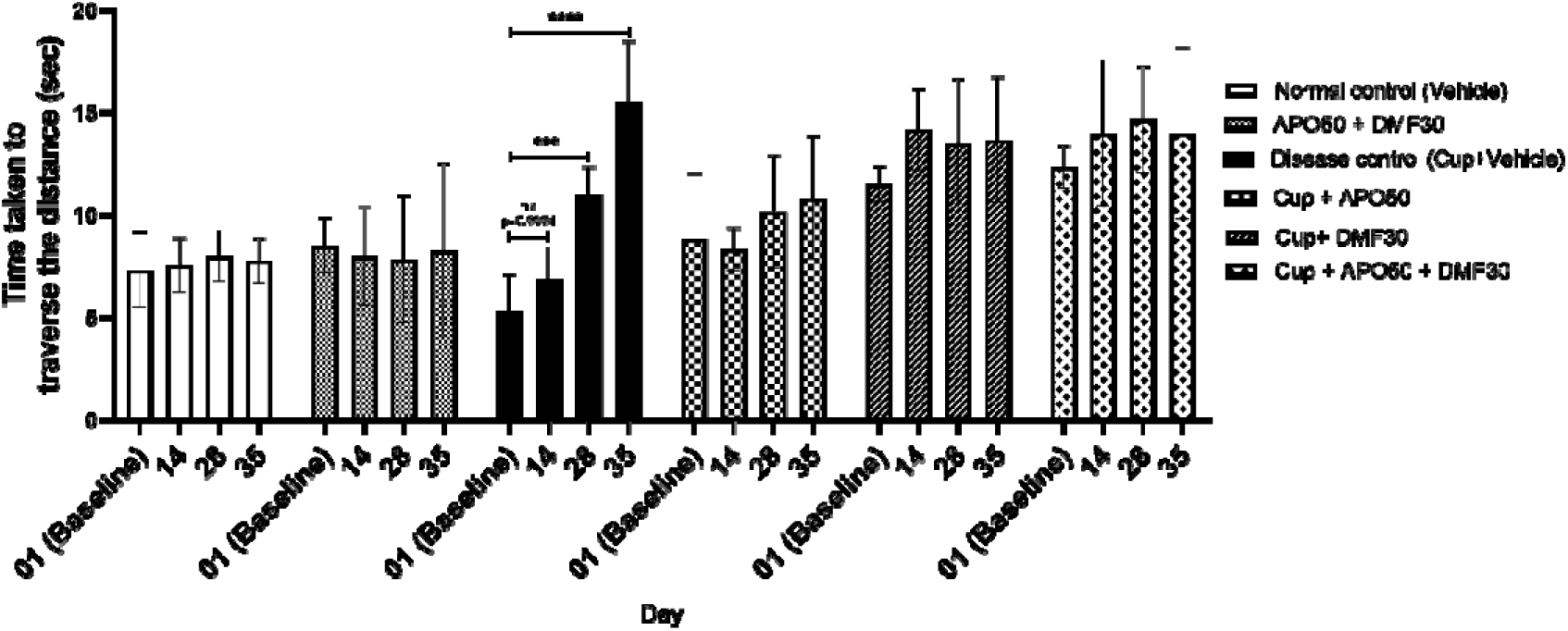
Narrow beam test. The time taken by the mice to traverse the distance (sec) was plotted on the y-axis against the day on the x-axis. The results are represented as mean ± s.d. ***, **** and ns indicates p<0.001, p<0.0001 and not significant, respectively.

**Fig. 5.**
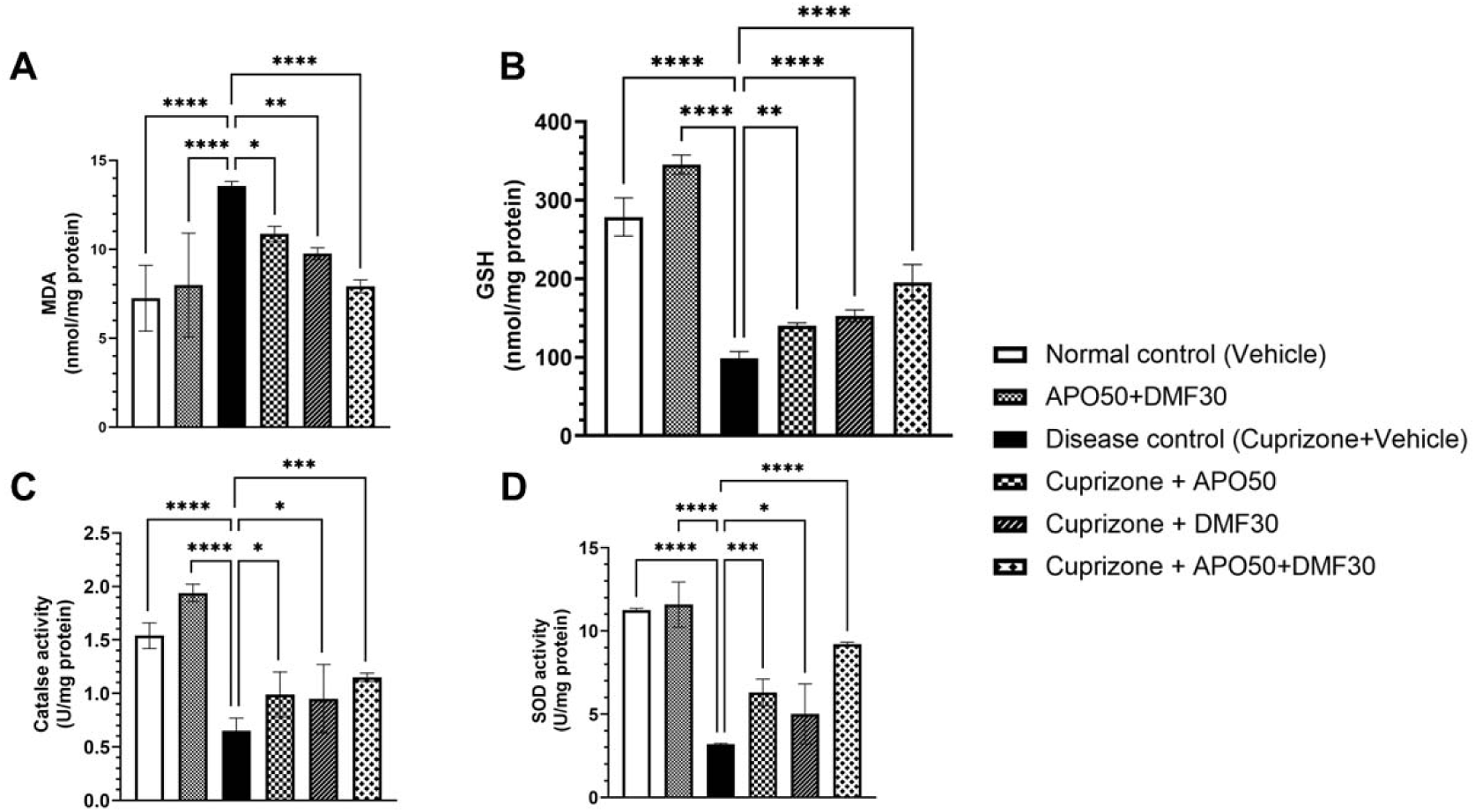
Levels of anti-oxidants and lipid peroxidation product. The levels of MDA (a), levels of GSH (b), levels of catalase (c), and levels of superoxide dismutase (d) against animal groups on the x-axis. The significance between the groups are represented by an asterisk (*), where *\*p*< 0.05, *\*\*p* < 0.01, *\*\*\*p* < 0.001, \*\*\*\**p*<0.0001.

### Effect of APO combination with DMF on glutathione

The analysis of data indicated a significant difference in the levels of GSH in CCO of brain (F (5, 24) = 186.3; p<0.0001). In CCO of the brain, the levels of GSH decreased significantly in disease control (98.52 ± 8.85 nmol/mg protein; p<0.0001) versus normal control (278.54 ± 24.22 nmol/mg protein; p<0.0001). MDA levels were significantly more in CCO of brain of cuprizone unfed, APO50 plus DMF30 treated mice (345.26 ± 12.22 nmol/mg protein; p<0.0001) compared to disease control. The levels of GSH significantly increased in the CCO of brain of CUP plus APO50 (140.45 ± 3.26 nmol/mg protein; p=0.0010), CUP plus DMF30 plus APO50 (195.25 ± 22.70 nmol/mg protein; p<0.0001), CUP plus DMF30 (152.85 ± 7.46 nmol/mg protein; p<0.0001) and with a 1.42-,1.55-, and 1.98-fold respectively, versus disease control (Fig.4B).

### Effect of Apocynin in combination with DMF on catalase activity

The analysis of data indicated a significant difference in the catalase activity of CCO of brain (F (5, 24) = 35.18, p<0.0001). In disease control, the CCO’s catalase activity significantly decreased (0.65 ± 0.12 U/mg protein; p<0.0001) compared to normal control (1.54 ± 0.12 U/mg protein), whereas in CCO of cuprizone unfed, APO50 plus DMF30 the catalase activity was significantly more (1.94 ± 0.08 U/mg protein) than disease control. Next, the CCO’s catalase activity was significant in CUP plus APO50 (0.99 ± 0.21 U/mg protein; p<0.05), CUP plus DMF30 (0.95 ± 0.32; U/mg protein; p<0.05), and CUP plus APO50 plus DMF30 (1.15 ± 0.04 U/mg protein; p<0.001) with 1.52-, 1.46-, and 1.76-fold changes versus disease control (Fig.4C).

### Effect of combination of APO and DMF on SOD activity

The analysis of CCO’s SOD activity indicated significant differences among the groups (F (5, 24) = 62.21, p<0.001). The CCO’s SOD activity significantly decreased in disease control (3.20 ± 0.05 U/mg protein; p<0.0001) versus normal control (Fig 4D). In CCO of cuprizone unfed, APO50 plus DMF30 the SOD activity was significantly more (11.58 ± 1.35 U/mg protein; p<0.0001) than disease control.

In CUP plus APO50, the SOD activity significantly increased (6.30 ± 0.80 U/mg protein; p=0.0002) versus disease control. The CCO’s SOD activity significantly increased in mice treated with DMF30 (5.01 ± 1.80 U/mg protein; p=0.0300) versus disease control. Next, DMF30 plus APO50 showed a significant increase in the brain (CCO) SOD activity (9.25 ± 0.10 U/mg protein; p<0.0001) with 2.89-fold changes, respectively versus disease control (Fig. 4D).

### Effect of apocynin combination with DMF on myelin in cuprizone-fed C57BL/6J mice

The analysis of scorings indicated significant differences among the groups (F_(5, 6)_ =19.07, p=0.0013). In normal control and cuprizone unfed APO50 plus DMF30 mice, the myelin structures in the corpus callosum (CC) area were intense, and significant (p<0.01) (Fig 6A and 6B) compared to disease control (CUP + Vehicle), where invisible staining in the CC area indicated severe myelin loss (Fig 6C).

**Fig. 6.**
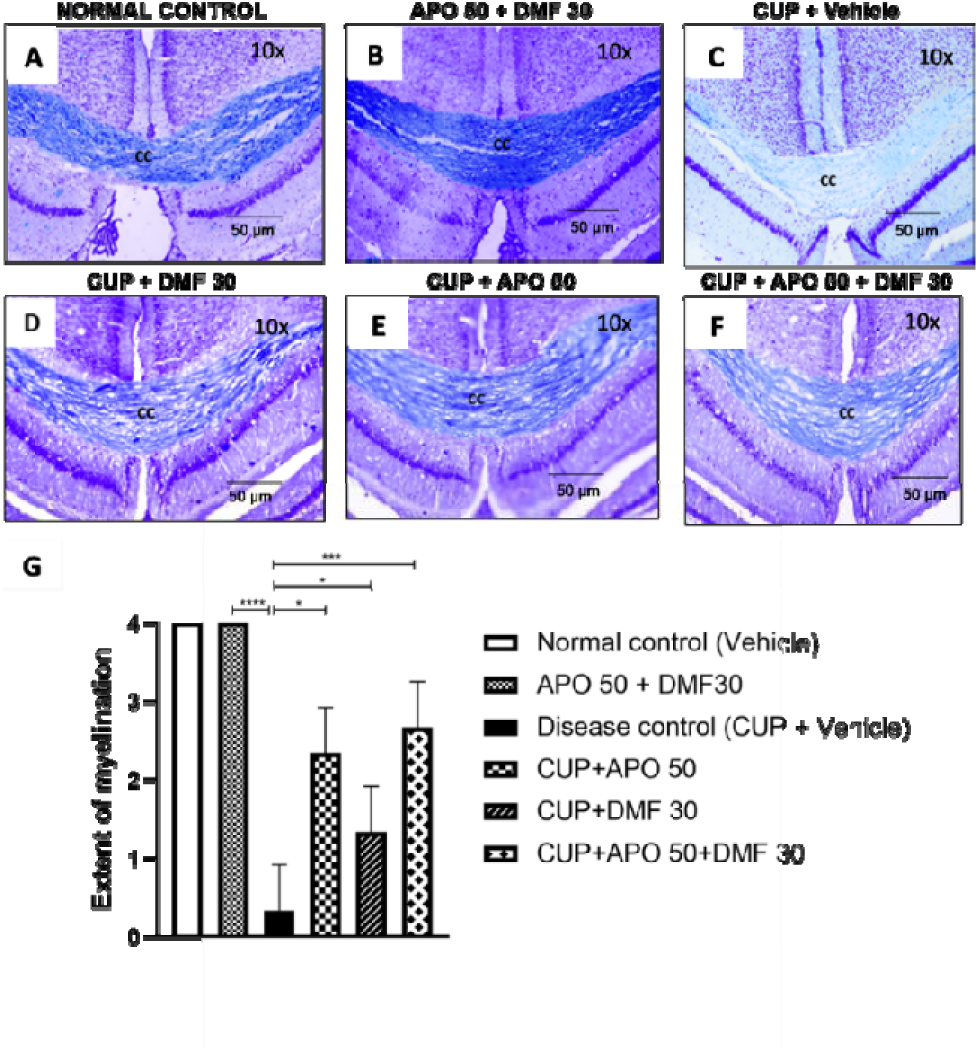
Modified luxol fast blue staining. Th e photomicrographs of coronal sections of mice brains stained with modified luxol fast blue are shown (animal groups indicated – right side). The mice brain sections were visualized under 10X magnifications and corpus callosum (CC) area was visualized (scale – 500 µm) for the staining of the myelin. The significance between the groups are represented by an asterisk (*), where *\*p*< 0.05, *\*\*p* < 0.01, *\*\*\*p* < 0.001, \*\*\*\**p*<0.0001.

In the treatment groups, a less intense, significant myelination (p= 0.0191) was observed in the CC area of CUP plus APO5O versus disease control (Fig. 6D). However, in the CC area of CUP+DMF30, the myelination was slightly preserved but insignificant (p=0.2363) (Fig. 6E) versus disease control. In fixed-dose combination group (CUP plus DMF30 plus APO50), the myelin was visible (6F) and significant in the CC area of mice brains (p=0.092) versus disease control.

## Discussion

Herein, we showed the effects of a fixed-dose of APO combination with a fixed-dose of DMF on motor coordination and long-term recognition memory in a cuprizone-induced mice model of demyelination. The mice exposed to cuprizone exhibit behavioral impairments, including abnormal motor coordination and cognitive damage [30,31]. In our study, mice fed with cuprizone (disease control) showed deficits in MC and LRM.

Herein, it was highly evident in disease control (CUP-fed), the latency to travel across the beam increased gradually indicating that CUP affected the motor coordination. A fixed-dose of either DMF or APO preserved motor coordination only moderately. A low dose of DMF (60 mg/kg) also moderately preserved MC in sodium nitroprusside-induced oxidative stress model [32]. However, in our study the treatment of combination of fixed-dose of APO plus DMF in CUP-fed mice preserved the latency and thereby MC, compared to their baseline.

CUP also affected the RI and DI, and treatment with fixed-dose of either DMF or APO was only moderate and not significant in preserving RI and DI compared to baseline. DMF enhanced cognitive performance in EAE mice [33] and apocynin was shown to increase learning and memory in various models of neurodegenerative diseases [34,35]. However, the treatment of combination of fixed-dose of APO plus DMF in CUP-fed mice preserved RI and DI compared to their baseline.

Next, we analyzed the levels of biochemical parameters of interest. It is well-known that CUP induces oxidative stress (OS), which also plays a significant role in MS pathogenesis [36]. Hence, we analyzed the levels of MDA, one of the markers involved in free radical-induced macromolecular damages in the brain [37,38]. Exposure to cuprizone has increased the levels of MDA; treatment with either DMF or APO, or their combination decreased the levels of MDA in the mice brains.

Out of other brain cells, oligodendrocytes require more metabolic supply to form myelin sheaths [39]. Endogenous antioxidants such as SOD and GSH are vital for cells to counteract oxidants [40].

DMF upregulates anti-oxidants by increasing mitochondrial membrane and cellular redox potential in a concentration-dependent manner [41]. Treatment of CUP-fed mice with DMF30 increased the brain levels of GSH in mice brains suggesting a protective action against oxidative stress [42,43]. Similarly, the treatment with APO50 elevated the levels of GSH in the CUP-fed mice owing to the inhibition of NOX [44]. Herein, the combination of APO plus DMF in CUP-fed mice elevated the levels of brain GSH versus monotherapy. Overall, DMF aided in the increased GSH levels; and APO protected the GSH levels through NOX scavenging. These effects on GSH production might be due to their differential mechanism, where DMF is one of the potent Nrf2 activators, and APO is a strong NOX inhibitor.

Nrf2 (Nuclear factor – erythroid factor 2-related factor 2) is a critical transcription factor and regulator of numerous cytoprotective genes [45]. Nrf2 is a cytosolic enzyme rapidly degraded by the ubiquitin-proteasome system. KEAP1 (Kelch-like ECH-associated protein 1) controls the stability and accumulation of Nrf2 by binding with the N-terminal Neh2 domain of Nrf2 [46]. In normal unstressed conditions, the cytoplasmic levels of Nrf2 are less; however, exposure to electrophilic chemicals or oxidative stress induces modification of reactive cysteine residues in KEAP1 (of Nrf2-KEAP1 complex), which stabilizes and elevate the cytosolic levels of Nrf2 [47]. Cytosolic Nrf2 translocates to the nucleus (nuclear translocation), binds to antioxidant response elements (ARE) of nuclear DNA, and transcribes mRNAs related to antioxidant proteins, including the enzymes for the synthesis of GSH [48]. Studies reported that DMF acts by depleting intracellular GSH and thiols modification on the Nrf2 inhibitor protein Keap1, which results in the stabilization of Nrf2 protein and increased expression of cytoprotective Nrf2 target genes [49]. Several *in vitro* cell-based studies reported that DMF, an Nrf2 activator found to increase the levels of GSH in glial cells [50,51]. Treatment with DMF also increased brain GSH in the mice models of cuprizone and EAE-induced demyelination [52].

Given cuprizone induces oxidative stress, one way through NOXs is by generating ROS [53]. Thus, inhibition of NOX is also essential to maintain the cellular levels of GSH. In the case of NOX, it is one of the primary sources of cellular ROS and is produced even under normal physiological conditions [54]. NOXs are found in the plasma membrane and in the membranes of phagosomes (of neutrophils) to engulf microorganisms [55]. However, it is activated to assemble in the membranes during a respiratory burst (i.e., oxidative stress). NOX catalyzes the production of a superoxide free radical by transferring one electron to oxygen from NADPH [56].

In our study, APO (NOX inhibitor) showed elevated levels of GSH in mice brains. This response might be due to the utilization of NADPH by glutathione reductase (due to inhibition of NOX) to convert oxidized glutathione (GSSG) to reduced GSH [57], thus protecting against oxidative stress. Evidence from studies found that inhibition of NADPH oxidase by APO promoted an anti-inflammatory effect during neuroinflammation [58] and decreased EAE-induced white matter damage in mice [17].

In addition, we also determined the activity of catalase and SOD in mice brains. The mice exposed to cuprizone for five weeks showed reduced brain catalase and total SOD activity. The combination therapy (DMF plus APO) has shown increased brain catalase and SOD activity. In several studies, DMF and APO have elevated the levels of SOD and catalase in mice brains with improvement in cognition [59,60].

The combined effects of DMF and APO also showed a positive association with the microscopic observation of myelin of the corpus callosum of mice, one of the most observed areas of white matter injury [61]. The demyelination in CC areas of the brain is one of the visible areas with white matter degradation, which can be well correlated with motor function impairment and cognition [62].

A study showed that the CUP-fed mice treated with DMF significantly protected myelin thickness and internodal length of axons in the CC area of brains [63]. Demyelination occurs maximum (peak injury) by the 4th and 5th week of CUP exposure [64]. Herein, end of the study (5th week), the LFB staining of mice brain sections indicated severe myelin loss in the corpus callosum of disease control. However, staining of brain sections isolated from mice treated with DMF plus APO in CUP-fed mice has shown more intense staining (presence of myelin) in the CC area than in other treatment groups.

## Conclusions

Therefore, DMF and APO combination might have protected myelin and preserved long-term recognition memory and motor coordination by elevating anti-oxidants in cuprizone-induced mice model of demyelination.

## List of abbreviations

MS: Motor coordination
LRM: Long-term recognition memory
MS: Multiple sclerosis
APO: Apocynin
DMF: Dimethyl fumarate
RRMS: Relapsing and remitting multiple sclerosis
OS: Oxidative stress
NADPH: Nicotinamide adenine dinucleotide phosphate
NOX: NADPH Oxidase
TBA: Thiobarbituric acid
TCEP: Tris (2-carboxyethyl)phosphine hydrochloride
ABD-F: 4-aminosulfonyl-7fluoro-2,1,3-benzoxadiazole
HPLC: High-performance liquid chromatography
GSH: Glutathione
MDA: Malondialdehyde
SOD: Superoxide dismutase
TET: Total exploration time
RI: Recognition index
DI: Discrimination index

## Declarations

### Ethical approval

The study protocol was approved by the Institutional Animal Ethics Committee (158/PO/ReBi/Re-L/99/CPCSEA), PSG Institute of Medical Sciences, Coimbatore, Tamil Nadu 641004, India. The experiments were carried out as per the guidelines of the Committee for the Purpose of Control and Supervision of Experiments on Animals (CPSCEA), New Delhi, India.

### Funding

Gurubarath Mani are supported through a post-graduate research fund, PSG College of Pharmacy, Peelamedu, Coimbatore, Tamil Nadu, India.

### Author’s contribution

The study was conceptualized and designed by Karthik Dhananjayan; the experiments were conducted, and data was analyzed and interpreted by Mani Gurubarath, and Karthik Dhananjayan; The first draft of the manuscript was written by Mani Gurubarath; further, the manuscript was modified and edited by Karthik Dhananjayan. The authors proofread AND approved the final manuscript.

## Acknowledgments

Authors acknowledge PSG Institutions, Peelamedu, Coimbatore, Tamil Nadu, India, for their generous and continuous support during this study.

## Notes

### Competing Interest Statement

The authors have declared no competing interest.

