## Supplemental files for "Multiple sclerosis: Effects of fixed-dose combination of dimethyl fumarate and NADPH oxidase inhibitor on oxidative stress markers and neurobehavioral activity in mice model of cuprizone-induced demyelination"

| **Content** | Page no |
| --- | --- |
| Supporting information. 1. Narrow beam test apparatus | 3 |
| Supporting information. 2. Bradford assay | 3 |
| Supporting information.3. Chromatogram of thiols | 4 |

**Supporting information. 1. Narrow beam test apparatus**

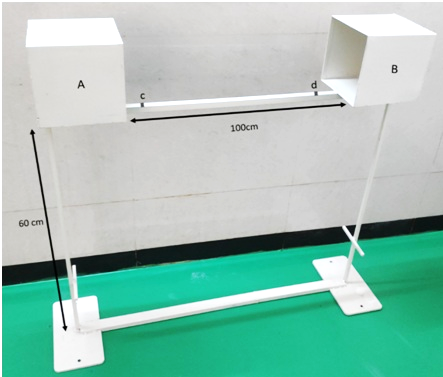

**Supporting information. 2. Bradford assay**

A serial dilution of 0, 10, 20, 40, 60, 80, and 100 ug/ml of BSA with a total volume of 20 µL in triplicates was added to the 96 micro-well titer plate. To each well, 200 µL Bradford reagent was added and read at 595nm. For the tissue sample, 20 µL of the brain supernatant was added in triplicates to the well, then 200 µL of Bradford reagent was added and read at 595nm.

**Supplementary information. 3. Chromatogram of thiols**

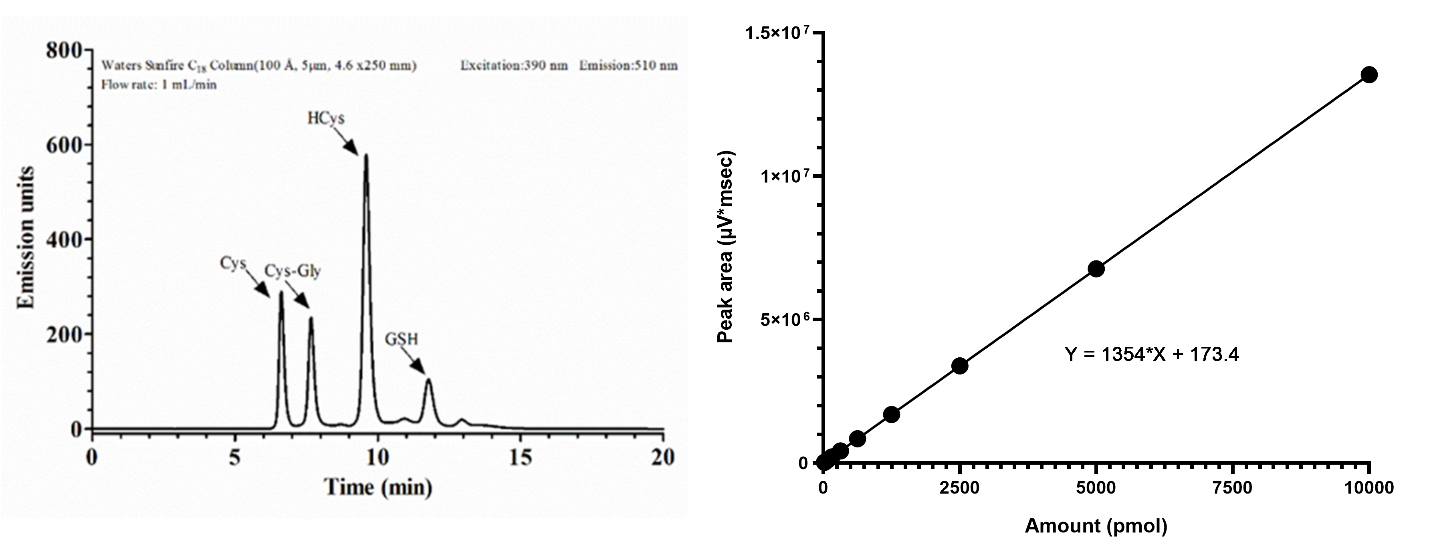

**Supporting information. 3. Chromatogram of thiols.** The retention time of 156 pmol of thiols. Cys: 6.52 min, Cys-Gly: 7.45 min, HCys: 9.55 min, and GSH: 11.92 min. In this study, the thiol of interest is GSH, and hence we quantified only the levels of GSH from their peak area after adjusting with dilution factor. The standard curve with a concentration on the x-axis and peak area on the y-axis. The linearity equation (y= mx + c) is also shown.
